# Novel design of imputation-enabled SNP arrays for breeding and research applications supporting multi-species hybridisation

**DOI:** 10.1101/2021.08.03.454059

**Authors:** G Keeble-Gagnère, R Pasam, KL Forrest, D Wong, H Robinson, J Godoy, A Rattey, D Moody, D Mullan, T Walmsley, HD Daetwyler, J Tibbits, MJ Hayden

## Abstract

Array-based SNP genotyping platforms have low genotype error and missing data rates compared to genotyping-by-sequencing technologies. However, design decisions used to create array-based SNP genotyping assays for both research and breeding applications are critical to their success. We describe a novel approach applicable to any animal or plant species for the design of cost-effective imputation-enabled SNP genotyping arrays with broad utility and demonstrate its application through the development of the Infinium Wheat Barley 40K SNP array. We show the approach delivers high-quality and high-resolution data for wheat and barley, including when samples are jointly hybridised. The new array aims to maximally capture haplotypic diversity in globally diverse wheat and barley germplasm while minimising ascertainment bias. Comprising mostly biallelic markers designed to be species-specific and single-copy, it permits highly accurate imputation in diverse germplasm to improve statistical power for GWAS and genomic selection. The SNP content captures tetraploid wheat (A- and B-genome) and *Ae. tauschii* (D-genome) diversity and delineates synthetic and tetraploid wheat from other wheats, as well as tetraploid species and subgroups. The content includes SNP tagging key trait loci in wheat and barley and that directly connect to other genotyping platforms and legacy datasets. The utility of the array is enhanced through the web-based tool *Pretzel* (https://plantinformatics.io/) which enables the array’s content to be visualised and interrogated interactively in the context of numerous genetic and genomic resources to more seamlessly connect research and breeding. The array is available for use by the international wheat and barley community.

**Short summary:** Designing SNP genotyping arrays for closely related species with broad applicability in both research and breeding is challenging. Here we describe a novel generic approach to select SNP content for such arrays and demonstrate its utility in wheat and barley to:

- capture haplotypic diversity while minimising ascertainment bias;
- accurately impute to high SNP density in diverse germplasm;
- generate high-quality high-resolution genotypic data; and
- jointly hybridise samples to the same bead chip array.

## Introduction

High-density genotyping arrays that simultaneously interrogate thousands of single nucleotide polymorphisms (SNP) have proven a powerful tool in genetic studies. The first generation of these have been widely used in wheat and barley for various applications including genome-wide association studies (GWAS), characterization of genetic resources, marker-assisted breeding and genomic selection (Pasam *et al*. 2017, Joukhadar *et al*. 2017, Balfourier *et al*. 2019). Continued advances in genome assembly and genotyping technologies present powerful new opportunities to continue the integration of genomics information into operational plant breeding systems and extend the potential of more academic research applications; e.g. studying genomic patterns of diversity, inferring ancestral relationships between individuals in populations and studying marker-trait associations in mapping experiments.

The publication of chromosome-scale genome assemblies are becoming available for more and more species and this availability is expected to accelerate with international projects such as the Earth BioGenome project (https://www.earthbiogenome.org/) which aims to sequence, catalog and characterize the genomes of all of the earth’s eukaryotic biodiversity over the next ten years. High quality assemblies are already available in cereal crop species such as barley (Mascher *et al.* 2017, Monat *et al.* 2019), emmer wheat (Avni *et al*. 2017), durum wheat (Maccaferri *et al.* 2019) and bread wheat (IWGSC 2018), as well as for the diploid ancestors of wheat (Luo *et al*. 2017, Ling *et al*. 2018). These assemblies have accelerated SNP discovery and our understanding of the breeding history of wheat and patterns of genome-wide linkage disequilibrium (LD) in different germplasm pools. For example, He *et al*. (2019) used an exome capture array in 890 globally diverse hexaploid and tetraploid wheat accessions to discover 7.3M varietal SNP and investigate the role of wild relative introgressions in shaping wheat improvement and environmental adaptation. Pont *et al*. (2019) exome sequenced a worldwide panel of 487 accessions selected from across the geographical range of the wheat species complex to explore how 10,000 years of hybridisation, selection, adaptation and plant breeding has shaped the genetic makeup of modern bread wheats. Similarly, Mascher *et al*. (2019) discovered almost 15M varietal SNP from exome sequence generated for 96 two-row spring and winter barley accessions, a subset of which was used to investigate the extent and partitioning of molecular variation within and between the two groups.

While SNP discovery using whole genome sequence data is currently limited to a relatively small number of wheat and barley accessions, this situation is expected to rapidly change as sequencing costs continue to decrease. For example, Lai *et al*. (2015) and Montenegro *et al*. (2017) used whole genome sequence data from 16 and 18 bread wheat accessions to identify more than 4M and 36M SNP on group 7 chromosomes and at the whole genome level, respectively. The more recent publication of whole genome sequence assemblies for 14 modern bread wheat varieties from global breeding programs (Walkowiak *et al*. 2020) provides additional new resources for *de novo* whole genome SNP discovery and to investigate structural variation within the wheat genome. In barley, Hill *et al*. (2020) used a combination of data sources including low coverage whole genome sequence of 632 genotypes representing major global barley breeding programs to investigate genomic selection signatures of breeding in modern varieties.

Increasing genomic resources and increased understanding of global and local population structure (Joukhadar *et al*. 2017) is enabling a shift from high to lower-density genotyping assays as a basis for undertaking genetic analyses for trait dissection and mapping. Where high-density data is still required, imputation can be effective to accurately infer higher marker density. Imputation uses statistical approaches to fill missing genotype data and increase low-density genotype data to genome-wide high-density data (Money *et al.* 2015). Imputation has been shown to increase power for the detection of marker-trait associations in GWAS (Jordan *et al*. 2015, Fikere e*t al.* 2020) and genomic selection (Nyine *et al.* 2019). Currently, hybridisation-based SNP arrays are better suited for imputation, compared to genotyping-by-sequencing (GBS) approaches, due to their lower missing data rates and higher genotype calling accuracies (Rasheed *et al*. 2017, Elbasyoni *et al.* 2018).

To date, several hybridisation-based SNP genotyping arrays providing genome-wide coverage have been developed for wheat and barley. Cavanagh *et al*. (2013) developed an Illumina iSelect array that genotyped 9,000 SNP. The same technology was used a year later to design an array that assayed 90,000 SNP (Wang *et al*. 2014), which was subsequently used to derive a breeder-oriented Infinium 15K array (Soleimani *et al*. 2020). Winfield *et al*. (2016) reported an Affymetrix Axiom 820K SNP array, which was also subsequently used to derive an Axiom 35K Wheat Breeders’ array that targeted applications in elite wheat germplasm (Allen *et al*. 2015). These genotyping arrays were largely based on genome sequence fragments from early Roche 454 and Illumina assemblies, or from exome capture sequence, and were generally enriched for gene-associated SNP. More recently, Rimbert *et al*. (2018) reported an Axiom 280K SNP array based on content derived from the intergenic fraction of the wheat genome, which to date has been poorly exploited for SNP, while Sun *et al*. (2020) described an Axiom 660K array based on genome-specific markers from hexaploid and tetraploid wheat, emmer wheat and *Ae. tauschii*. In barley, two Infinium iSelect genotyping arrays comprising 9K and 50K SNP have been reported (Comadran *et al*. 2012, Bayer *et al*. 2019).

While SNP genotyping arrays provide robust allele calling with high call rates and fast sample turn around (typically about 3 days), they have high set up costs. The latter has presented significant challenges for the development of SNP arrays that can comprehensively serve both research and breeding applications; researchers have traditionally preferred high SNP density (which creates a high genotyping cost per sample but low cost per datapoint), while breeders typically only want a minimally sufficient marker density. This challenge drove us to develop a general approach to SNP array design that specifically takes into consideration the need for low-cost genotyping across a wide range of research and breeding applications, with the aim to seamlessly connect research to breeding.

Here, we present the design methodology and an example of its implementation in the Infinium Wheat Barley 40K SNP array Version 1.0, a new and highly optimised genotyping platform containing 25,363 wheat-specific and 14,261 barley-specific SNP, the vast majority of which behave as easily scored, single-copy biallelic markers. The SNP content was carefully selected to enable accurate imputation to high SNP density in globally diverse wheat and barley germplasm, as well as within the more restricted germplasm pools of breeding programs. The array is well connected to markers on other commonly used SNP arrays, as well as to many existing genomic resources, and provides high utility in research and breeding from germplasm resource characterisation, GWAS and genetic mapping to tracking introgressions from different sources, marker-assisted breeding and genomic selection. In addition, the SNP have been selected to enable joint hybridisation of wheat and barley samples in the same assay, potentially halving costs for large scale deployment. The array is available for use by the international wheat and barley community and is supported by the web-tool *Pretzel* (Keeble-Gagnère *et al*. 2019, https://plantinformatics.io/).

## Materials and Methods

### Germplasm and genomic resources

SNP genotypes for 1,041 exome sequenced bread wheat accessions were used to select content for the Infinium Wheat Barley 40K SNP array. The accessions included 790 previously reported in He *et al*. (2019) to capture global wheat diversity, an additional 149 accessions selected from the global collection contained in the associated VCF file (http://wheatgenomics.plantpath.ksu.edu/1000EC/) to expand the diversity captured and 102 historical breeding lines from the InterGrain commercial wheat breeding program (https://www.intergrain.com). The first two sets of accessions maximally captured genetic diversity among 6,087 globally diverse wheat accessions comprising landraces, varieties, synthetic derivatives and novel trait donor lines (He *et al.* 2019). The additional 149 accessions were selected to capture genetic diversity within synthetic derivative germplasm derived from crossing 100 primary synthetics (derived from interspecific hybridisation of durum wheat with *Ae. tauschii*) to three Australian varieties: Yitpi, Annuello and Correll (Ogbonnaya *et al*. 2007). The latter two sets of accessions were exome capture sequenced as described in He *et al*. (2019). SNP discovery was performed using the first two sets of accessions and the resulting SNP list was used to call SNP genotypes across all accessions.

The Infinium 90K wheat SNP genotypes reported in Maccaferri *et al*. (2019) for a globally diverse tetraploid wheat collection of 1,856 accessions comprising wild emmer (*Triticum turgidum* ssp. *dicoccoides*), domesticated emmer (*T. turgidum* ssp. *dicoccocum*) and *T. turgidum* genotypes including durum landraces and cultivars were used to select tetraploid wheat specific SNP.

A georeferenced landrace collection of 267 exome sequenced barley accessions, including 2- and 6-rowed *Hordeum vulgare* landraces as well as *Hordeum spontaneum* (Russell *et al*. 2016), and 117 whole genome sequenced accessions representing historical breeding lines from the InterGrain commercial barley breeding program were used to select content for the SNP array.

### SNP discovery

In wheat, SNP discovery and genotype calling were performed as described in He *et al.* (2019), against IWGSC RefSeq v1.0 (IWGSC 2018). After filtering for >40% call rate and >1% minor allele frequency (MAF), 2.04M SNP were used for LD analysis. To filter for nucleotide variation originating from *Ae. tauschii*, D-genome-specific SNP that had a MAF >0.1 in the synthetic derivative wheat and MAF <0.1 in the globally diverse wheat collection were identified. In addition, the top 2% of D-genome SNP that showed differential allele frequencies between these two groups based on Fst values (Weir and Cockerham 1984) were selected. From these two SNP sets, SNP uniformly distributed across the D-genome were selected for inclusion as SNP content.

In barley, SNP discovery was performed as described in He *et al*. (2019) using the exome sequence data published in Russell *et al.* (2016), against Morex v1.0 (Mascher *et al.* 2017). Following removal of *Hordeum spontaneum-*like accessions based on PCA clustering (which left 157 *Hordeum vulgare*-like accessions), the resulting SNP list was used to call SNP genotypes in the 120 InterGrain historical breeding lines. After filtering for >40% call rate and >5% MAF (a higher cut-off was used in barley due to the smaller reference population), 932,098 SNP were used for LD analysis.

### Linkage disequilibrium analysis

LD analysis for the filtered SNP was performed using PLINK (Purcell *et al.* 2007) at the chromosome level within each species with a maximum window size of 2 Mb; i.e. all the SNP in a tag SNP set had to be within a 2 Mb window. The squared correlation coefficient (*r*^2^) based on the allele frequency in the global barley or wheat diversity panel (excluding the synthetic derivatives) between two SNP was considered as a measure of LD.

### Choice of SNP probe designs

To maximise the number of SNP assayed for a given number of probes on the bead chip array, A/T and C/G variants (Infinium Type I SNP which require two probes) were avoided. To maximise SNP scorability and genotype calling accuracy, polymorphism underlying the 50-mer oligonucleotide SNP probe sequences was also avoided as they are known to cause shifts in SNP cluster position (Wang *et al*. 2014). For tSNPs, the probe sequences were required to align uniquely to the target genome and not align to the other genome; i.e. a wheat SNP probe had to align uniquely to the wheat genome and not to the barley genome, and vice versa. Finally, an Illumina Design Tool score of ≥0.6 was required for a probe to be included as array content. A relaxed set of criteria was also used (to tag SNP sets otherwise missed) which allowed up to 3 alignments to the target genome.

### Selection of tagging SNP (tSNP) for imputation

A custom algorithm was used to select tSNP tagging LD blocks in each of the global collections and to facilitate imputation from the density of the SNP array. In brief, for each chromosome the algorithm iteratively selected the most informative tSNP passing all filters (based its *r*^2^ value from the LD analysis), removed all SNPs linked to the selected tSNP from the remaining list of SNPs, as well as all SNP linked to any SNP in the selected tSNP set to avoid directly tagging any SNP at *r*^2^ ≥0.9 more than once, before repeating the process until a target number of tSNP was reached. This process ensured the set of tSNP selected was the minimum set required to tag the most SNPs at *r*^2^ ≥0.90. Specifically, for a given a set of SNPs *S* = {*s*_1_, *s*_2_, …} and function *r*^2^(*s*_*i*_, *s*_*j*_) defining the *Pearson correlation squared* ∀*s*_*i*_, *s*_*j*_ ∈ *S*, we defined the tSNP set for *s*_*i*_ at *q* to be:

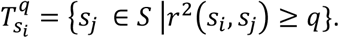

Rename the 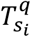 and define 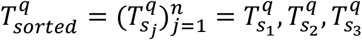, … where 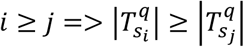.

In other words, 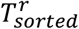 is an ordering of tSNP sets, monotonically decreasing in size. Let *F* ⊂ *S* be a subset of filtered SNPs. Define 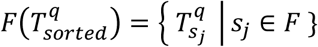.

We define 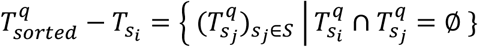, and *head*(*L*) to be the first element of the ordered sequence *L*.

The algorithm is then:

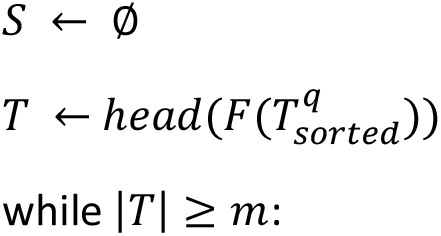
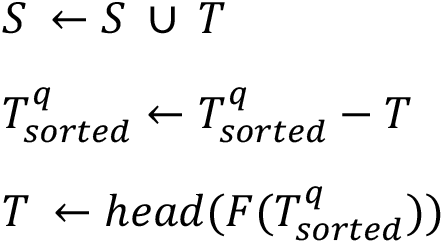

The above is applied with *q* = 0.9, *m* = 10 to define the imputation set *S*.

To guard against possible loss of imputation accuracy due to SNP assays failing to provide reliable genotypes calls, a level of redundancy was included in the tSNP sets for wheat and barley. Specifically, three tSNP were chosen when the number of SNP tagged was ≥50 and two tSNP were selected when the number of SNP tagged was ≥20. Single tSNP were included as array content when they tagged at least 10 SNP. Some tag SNP sets could not be tagged because no probe passed all filters; in this case we ran the algorithm on the remaining sets allowing SNP passing relaxed filters (up to 3 hits to target genome were allowed). In addition, tSNP were selected to tag genomic regions that had sparse SNP coverage but high LD; i.e. tagging <10 SNP within windows larger than 500Kb in wheat and 1Mb in barley. Finally, SNP were selected in regions still lacking SNP after the previous steps.

### Optimisation of SNP content

To ensure broad applicability of the SNP array in research and breeding, the content included SNP selected to specifically interlink germplasm resources such as the 19,778 domesticated barley accessions with GBS genotypes described in Milner *et al.* (2019). It also included SNP probes designed to interrogate published trait-linked markers in wheat and barley. Designs for these markers were based directly on published sequence or from alignment of published primers or flanking sequences and inference of the targeted nucleotide variation. For all trait-linked markers, the best probe design was selected based solely on the Illumina quality score. Due to difficulty for designing SNP probes targeting known alleles of phenology genes, we selected 293 exome SNPs around the genes reported in Shi *et al*. (2019).

### Imputation

The wheat and barley global diversity sets were used as reference haplotypes for imputation. For wheat, accessions clustering with the synthetic derivatives in a PCA analysis were excluded. For barley, only samples with <20% missing data were used. In both species, missing data was filled in using Beagle (Browning *et al.* 2007) and phased with Eagle (Loh *et al*. 2016). In total, 868 and 155 wheat and barley lines were used as reference haplotypes.

In wheat, SNP coordinates were converted to IWGSC v2.0 pseudomolecules (https://urgi.versailles.inra.fr/download/iwgsc/IWGSC_RefSeq_Assemblies/v2.0/, Zhu *et al*. 2021) before imputation. After transfer into the v2.0 assembly, there were 18,521 SNP before imputation, with 630,058, 549,003 and 352,947 tagged at *r*^2^ ≥0.50, 0.70 and 0.90, respectively.

To assess the accuracy for imputation into globally diverse germplasm, 100-fold cross validation was performed. A random subset of 100 wheat (or 10 barley) lines had their true genotypes masked, leaving only the tSNP. The remaining lines were then used as the reference population with Minimac3 software (Das *et al*. 2016) to impute back the missing genotypes for three different target SNP sets: the set of SNP tagged at *r*^2^ ≥0.50, 0.70 and 0.90, respectively. The imputation accuracy for each line, measured as both correlation and concordance between the actual and imputed genotypes, was calculated from 100 repetitions of this process in each of wheat and barley. Correlation was measured as the Pearson *r*^2^ between SNP called in both genotypes being compared, while concordance was measured as the fraction of SNP in agreement between those called in both genotypes being compared.

### SNP assay and genotype calling

Samples were assayed following the protocol for Infinium XT bead chip technology (Illumina Ltd). SNP clustering and allele calling was performed using GenomeStudio Polyploid software (Illumina Ltd) using the Illumina-supplied wheat or barley SNP manifest file. The custom genotype calling pipeline described in Maccaferri *et al*. (2019) was also used.

## Results

### Overview of design approach

The central idea of the design concept is to exploit LD using the *r*^2^ measure to define sets of SNP that can be considered equivalent: for a given SNP, we define its tag SNP set as the set of SNP with *r*^2^ ≥ 0.9 (the set of SNP in this set are referred to as tSNP). This metric provides a measure of equivalence as well as a natural ranking of SNP by their informativeness, as defined by the size of the tSNP set to which they belong. We assume the relationship is symmetrical; i.e. if SNP A is in SNP B’s tSNP set, then SNP B should be in SNP A’s tSNP set. The original set of SNP is then filtered using technology and application-specific criteria (see Materials and Methods) while maintaining connectivity to SNP that fail the filters via the tSNP sets of SNP that pass the filters.

To design a genotyping array that has broad applicability in research and breeding, the SNP should be discovered in diverse germplasm to avoid ascertainment bias (since LD is population dependent) and with sufficient density to produce large tSNP sets. The latter helps ensure at least one SNP in a tSNP set will pass all the design filters in most instances. Here, we used a globally diverse set of barley landrace accessions and a globally diverse set of wheat accessions that included landraces, varieties, novel trait donor and historical breeding lines (Figure 1). For array designs focused only on breeding applications, SNP discovery should aim to capture the genetic diversity within the breeding germplasm pool.

**Figure 1.**
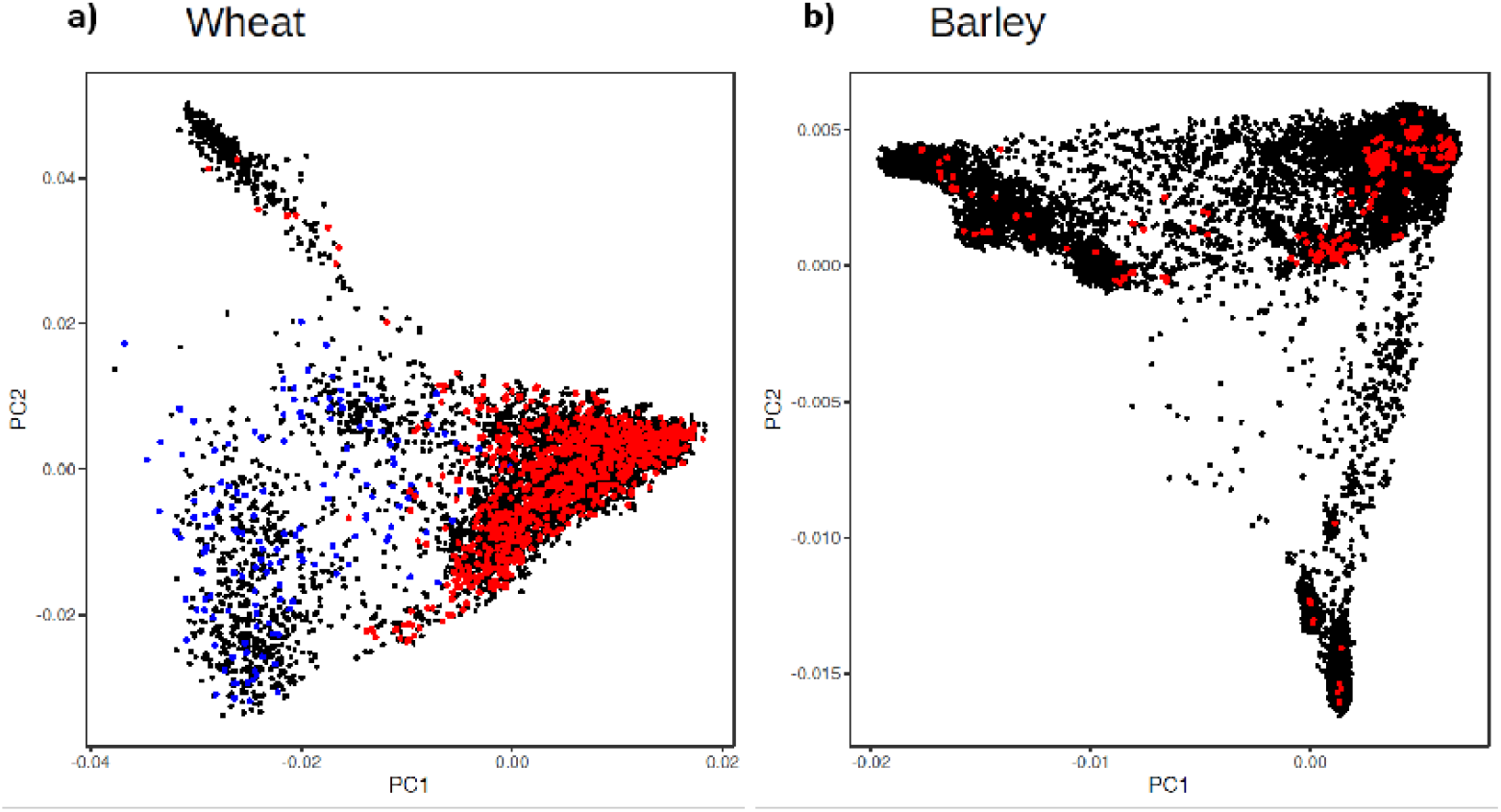
PCA plots showing genetic diversity of wheat and barley accessions used for SNP discovery. (**a**) 6,087 wheat accessions genotyped with the iSelect wheat 90K SNP array (Wang *et al*. 2014) (black), exome-sequenced accessions used for LD analysis (red) and synthetic derivative accessions capturing D-genome diversity (blue); and (**b**) 19,778 barley accessions genotyped with GBS (black), with exome-sequenced accessions used for LD analysis (red).

A novel selection algorithm (described in Materials and Methods) is then used to select SNP which maximise LD capture, while minimising the number of SNP assayed on the array, using only SNP that pass the design filters.

The design concept can be applied to any animal or plant species. In addition to this set of SNP, utility in research and breeding can be further enhanced by including context-relevant SNP such as trait-linked markers and markers that link germplasm resources across different genotyping technologies.

### SNP discovery and filtering

Filtering for a minimum minor allele frequency (MAF) of 1% and maximum missing rate of 60% using the 8,869,370 wheat SNP published in He *et al.* (2019) resulted in 2,037,434 high quality SNP for downstream analysis. Of these, 122,799 SNP had at least one array probe that passed all design filters. In barley, filtering of the 1,843,823 SNP identified from our processing of exome capture sequence from the accessions from Russel *et al*. (2016) for MAF >5% and missing rate <60% resulted in 932,098 high quality SNP for downstream analysis, of which 119,633 SNP had at least one array probe passing all filters. The filtered SNP matrices used in subsequent analysis are available at https://dataverse.harvard.edu/dataverse/WheatBarley40k_v1.

### LD analysis and selection of tagging SNP for imputation

Based on LD values of *r*^2^ ≥0.9, a total of 1.07M wheat and 413,508 barley high quality SNP were singletons; i.e. had no SNP within 1Mb up- and downstream with *r*^2^ ≥0.9. These SNP were either genuine singletons or categorised as singletons due to the absence of additional SNP within the surrounding 2Mb region. As singleton SNP can only be tagged directly, which is not feasible on a low-density array, these SNP were not considered further for inclusion on the array.

The custom selection algorithm grouped the 122,799 non-singleton wheat SNP passing all design filters into 11,076 tSNP tagging SNP sets containing ≥10 SNP within a 2 Mb window. These tSNP tagged 317,599, 538,326 and 652,476 SNP at *r*^2^ ≥0.9, 0.7 and 0.5, respectively. Of the 119,633 non-singleton barley SNP passing all filters, the selection algorithm identified 7,316 tSNP which tagged a total of 150,096, 294,659 and 390,844 SNP at *r*^2^ ≥0.9, 0.7 and 0.5, respectively. At the genome level, the rate of return per tSNP was surprising similar for wheat and barley and plateaued at about 15,000 tSNP at *r*^2^ ≥0.9 (Figure 2). However, the rate of return per tSNP varied at the chromosome level (Figure S1).

**Figure 2.**
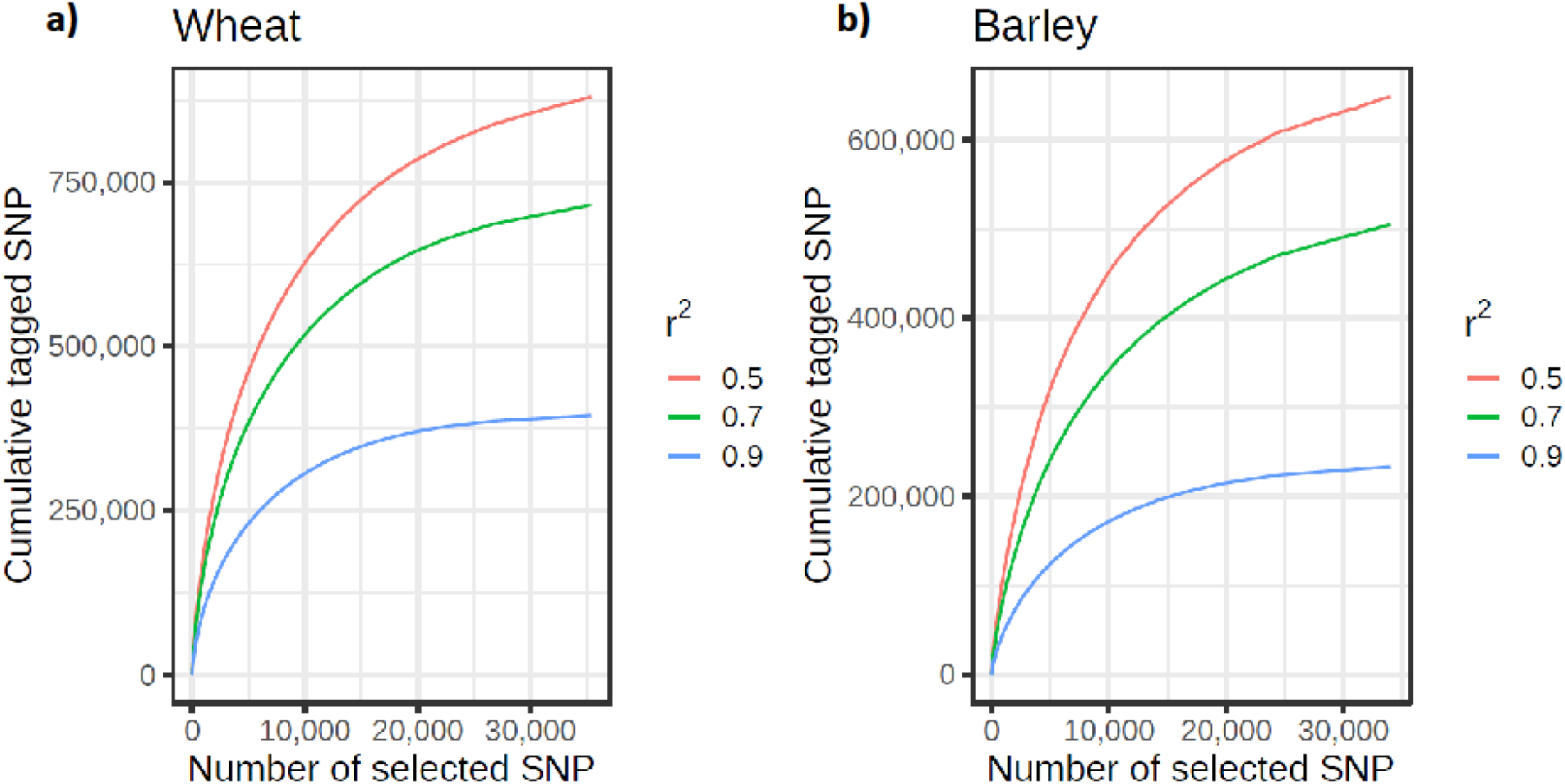
Cumulative number of SNP tagged by tSNP at *r*^2^ ≥0.9, 0.7 and 0.5 respectively in wheat and barley. The curves are shown until the first singleton SNP (at *r*^2^ ≥0.90) is reached

In total 21,012 wheat and 13,469 barley tSNP were included as content on the array. This tally includes redundant SNP selected to guard against possible loss of imputation accuracy due to SNP assays that might fail; SNP passing a relaxed set of filters (allowing up to 3 alignments to the target genome) and tagging SNP sets untaggable with the stricter filtered SNP; and SNP to tag genomic regions that had sparse SNP coverage but high LD; i.e. tagging <10 SNPs within windows larger than 500Kb in wheat and 1Mb in barley. The latter SNP are expected to support increased imputation density in these regions as higher density SNP datasets become available into the future. The wheat tSNP tagged a total of 394,034, 636,641 and 758,452 SNP at *r*^2^ ≥0.9, 0.70 and 0.50 respectively, while the barley tSNP tagged a total of 187,412, 361,012 and 471,645 SNP, respectively. Importantly the MAF distributions for the tSNP, tagged SNP and filtered SNP from the globally diverse wheat and barley collections closely matched one another, respectively (Figure 3).

**Figure 3.**
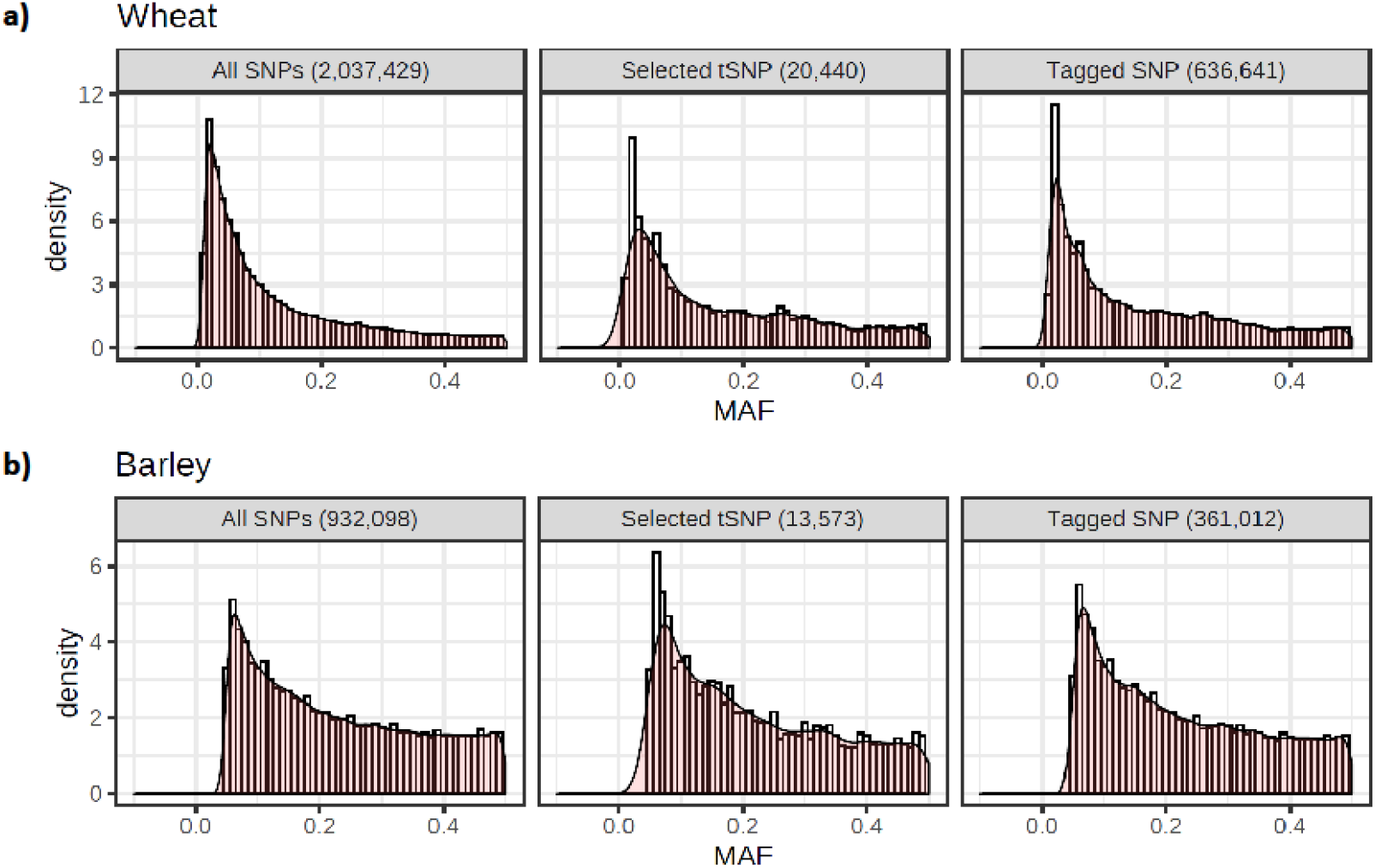
MAF distribution of all SNP used for LD analysis, selected tSNP and the set of SNP tagged by the tSNP at *r*^2^ ≥0.70 in the globally diverse wheat (n=790) and barley (n=157) collections

### Accuracy for imputing into globally diverse germplasm

The ability to impute from the tSNP on the array to the sets of SNP tagged at *r*^2^ ≥0.50, 0.70 and 0.90 respectively in globally diverse wheat and barley germplasm was assessed using 100-fold cross validation. Accuracy was determined from the correlation and concordance between the imputed and actual genotypes for each wheat or barley line averaged over the occurrences of that sample within the 100 iterations.

As expected, all metrics were highest when imputing to the set of SNP tagged at *r*^2^ ≥0.90 and lowest for those tagged at *r*^2^ ≥0.50 (Table 1). In wheat, only a small decrease in accuracy was observed for most accessions as the size of the tagged SNP set increased (i.e. *r*^2^ decreased), with reduced accuracy most evident in the bottom 50 accessions (Figure 4). For these accessions, the difference in accuracy (both correlation and concordance) between comparisons including and excluding heterozygous genotype calls was almost 10%, suggesting the possibility of high error rates in the heterozygous exome SNP calls for these accessions. 768 (88.5%) of the wheat accessions had accuracies ≥90% with the strictest correlation metric (which included heterozygous calls) for the set of SNP tagged at *r*^2^ ≥0.50. When comparing only non-heterozygous calls, the number of lines above this threshold rose to 866 (99.8%) (Figure 4).

**Table 1.**
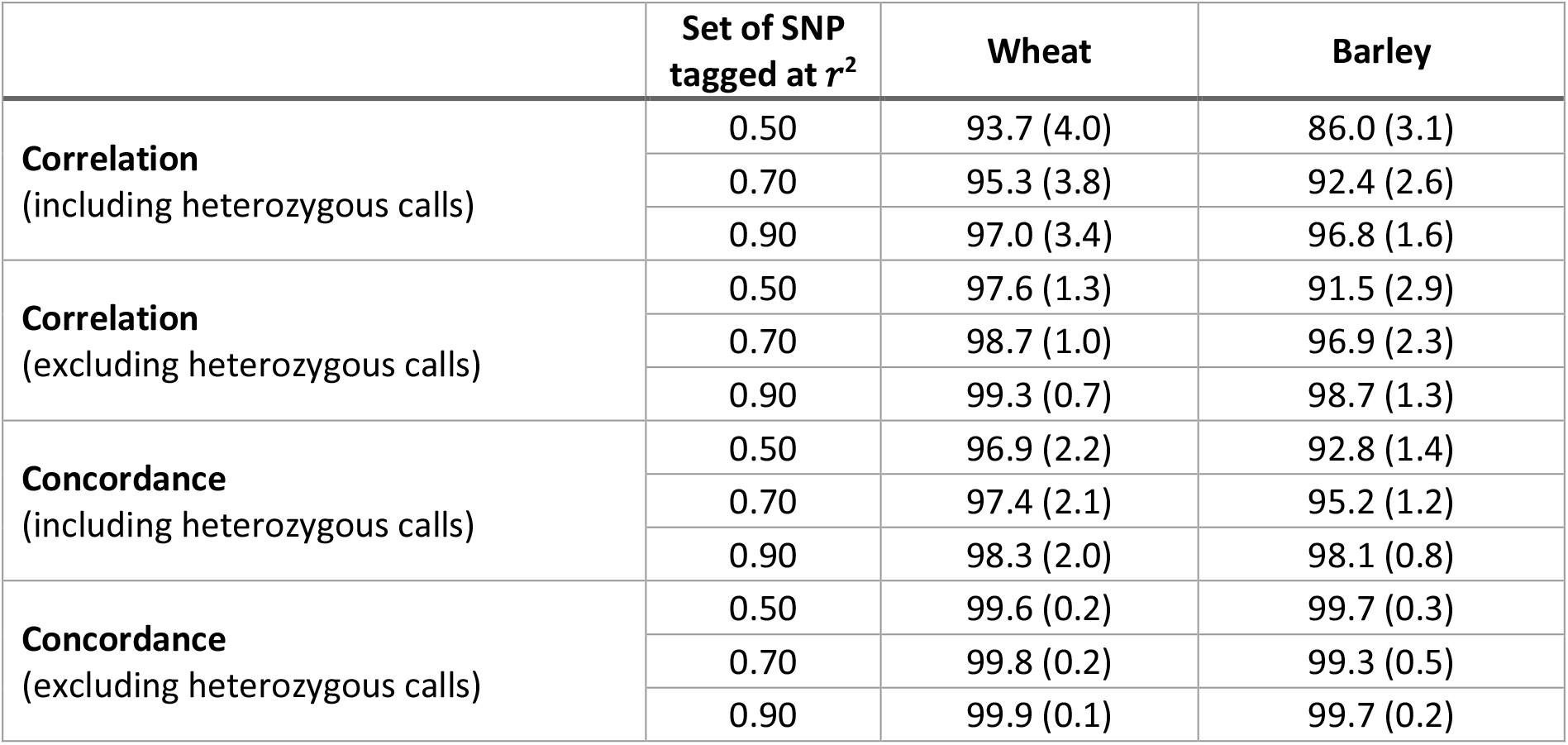
Accuracy for imputing from the tSNP on the array to the sets of SNP tagged at *r*^2^ ≥0.50, 0.70 and 0.90 respectively in wheat and barley. Correlation is the Pearson *r*^2^ between SNP called in both genotypes being compared. Concordance is the fraction of SNP in agreement between those called in both genotypes being compared. Standard deviations are shown in brackets

| | Set of SNP<br>tagged at $r^2$ | Wheat | Barley |
| --- | --- | --- | --- |
| **Correlation**<br>(including heterozygous calls) | 0.50 | 93.7 (4.0) | 86.0 (3.1) |
|  | 0.70 | 95.3 (3.8) | 92.4 (2.6) |
|  | 0.90 | 97.0 (3.4) | 96.8 (1.6) |
| **Correlation**<br>(excluding heterozygous calls) | 0.50 | 97.6 (1.3) | 91.5 (2.9) |
|  | 0.70 | 98.7 (1.0) | 96.9 (2.3) |
|  | 0.90 | 99.3 (0.7) | 98.7 (1.3) |
| **Concordance**<br>(including heterozygous calls) | 0.50 | 96.9 (2.2) | 92.8 (1.4) |
|  | 0.70 | 97.4 (2.1) | 95.2 (1.2) |
|  | 0.90 | 98.3 (2.0) | 98.1 (0.8) |
| **Concordance**<br>(excluding heterozygous calls) | 0.50 | 99.6 (0.2) | 99.7 (0.3) |
|  | 0.70 | 99.8 (0.2) | 99.3 (0.5) |
|  | 0.90 | 99.9 (0.1) | 99.7 (0.2) |

**Figure 4.**
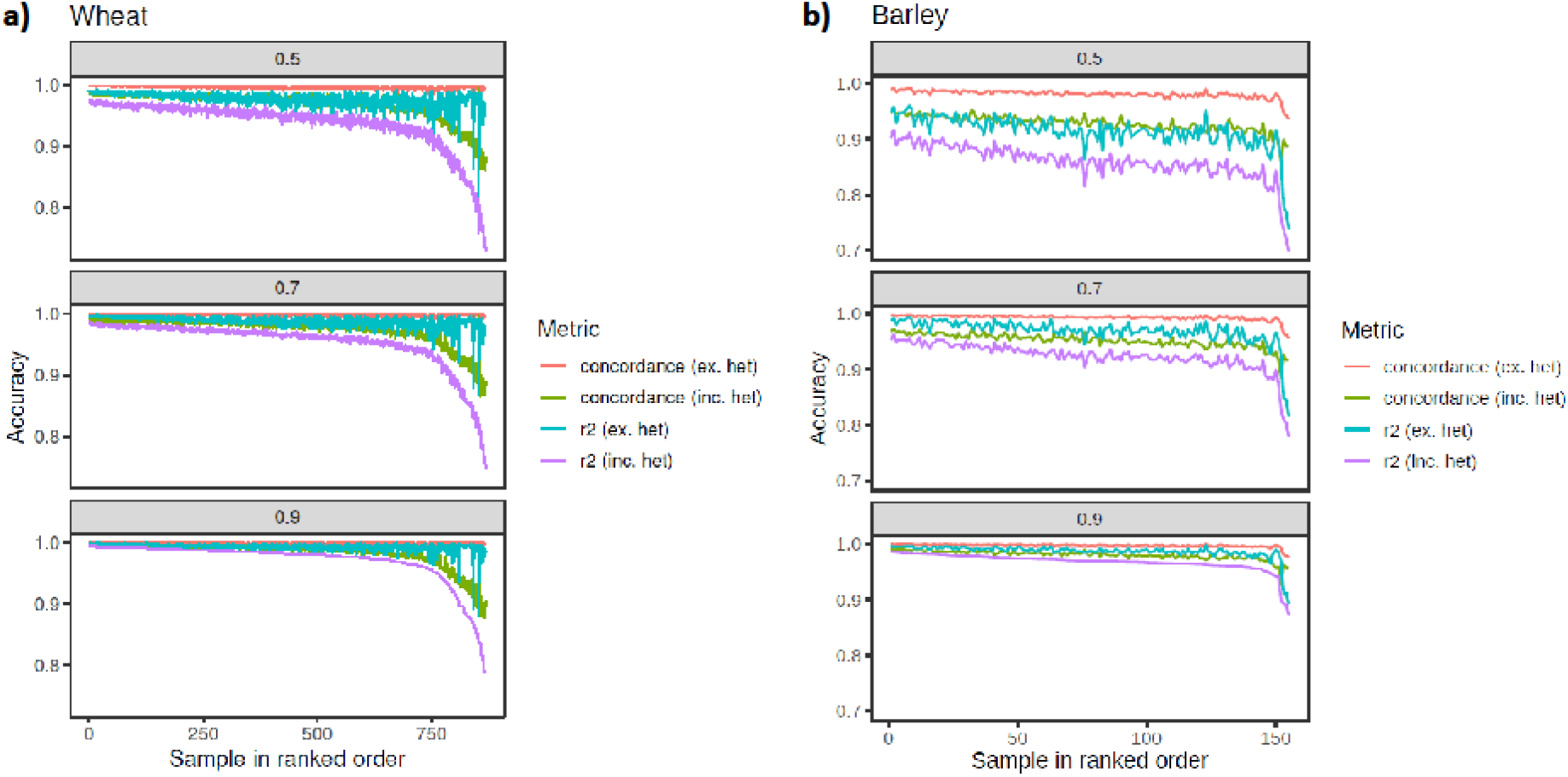
Imputation accuracy from the tSNP on the array to the set of SNP tagged at *r*^2^ ≥0.5, 0.7 and 0.9 respectively in wheat and barley. Metrics plotted are correlation *r*^2^ including heterozygous calls (orange line), *r*^2^ excluding heterozygous calls (cyan line), concordance including heterozygous calls (green line) and concordance excluding heterozygous calls (orange line). The accessions are ranked ordered based on the *r*^2^ including heterozygous calls

Reduced accuracy when imputing to higher tagged SNP numbers was more pronounced in barley. A difference of 10.8% (from 96.8% to 86%) was observed between the average correlation (which included heterozygous calls) for the set of SNP tagged at *r*^2^ ≥0.90, compared to those tagged at *r*^2^ ≥0.50 (Table 1). As observed in wheat, the inclusion of heterozygous calls reduced the accuracy, particularly when imputing to the set of SNP tagged at *r*^2^ ≥0.50, again suggesting possible erroneous heterozygous calls in the sequence genotypes (Figure 4). The reduced accuracies observed in barley compared to wheat are also likely partly due to the reduced size of reference haplotypes (155 versus 868). Accuracies in barley would likely improve if the reference haplotype set was expanded.

### Wheat-barley 40K SNP array content

The final array design comprised 34,481 imputation SNP and two additional categories of context-specific SNP (content summarised in Table 2, full details are in Table S1).

**Table 2.** SNP content of the Infinium Wheat Barley 40K SNP bead chip array

|  | **Wheat** | **Barley** | **Total** |
| --- | --- | --- | --- |
| Tagging SNP for imputation | 21,012 | 13,469 | 34,481 |
| Trait associated SNP | 427 | 178 | 605 |
| SNP linking germplasm resources | 3,924 | 614 | 4,538 |
| **Total number of SNP** | **25,363** | **14,261** | **39,624** |

The first context-specific category included 2,609 SNP from the Infinium wheat 90K SNP array (Wang *et al.* 2014) that were selected based on allele differentiation to tag tetraploid wheat (A- and B-genome) diversity and to clearly delineate tetraploid wheat from other types of wheat, as well as distinguish tetraploid species and subgroups from one another. The SNP comprised four classes: 1) differentiating SNP that represent the top 2% Fst values in Maccaferri *et al*. (2019) between the four subgroups of tetraploid species: wild emmer, domesticated emmer, domesticated wild emmer, durum landraces and durum cultivars; 2) subgroup-specific private SNP that showed a MAF ≥0.1 in one of the subgroups and were either monomorphic or showed a MAF <0.05 in the other subgroups; 3) subgroup-specific high MAF SNP that were present at ≥0.3 MAF in any one of the subgroups; and 4) neutral SNP that did not show any signatures of selection, were polymorphic in all subgroups and showed an overall MAF of ≥0.4. The ability of these SNP to reliably differentiate the tetraploid species subgroups as efficiently as the Infinium wheat 90K array is shown in Figure S2.

The second category included 1,206 exome SNP tagging *Ae. tauschii* (D-genome) diversity present in backcross synthetic derivatives that originated from crosses involving 100 primary synthetic parents, which were selected for phenotypic and genetic diversity among about 400 primary synthetics developed at CYMMIT and imported into Australia in 2001. Each of the 100 primary synthetic parents was derived from a different *Ae. tauschii* accession. The SNP were selected to provide high D-genome coverage, enriched density in highly recombining chromosomal regions and to clearly delineate bread wheat from other types of wheat, as well as tag diversity in synthetic wheat and their derivatives and *Ae. tauschii*. The SNP comprised two classes: 1) differentiating SNP that represent the top 2% Fst values between the global diversity wheat and synthetic derivative collections; and 2) D-genome diversity from *Ae. tauschii* that showed a MAF ≥0.1 in the synthetic derivative collection and MAF ≤0.1 in the global diversity wheat collection. The ability of these SNP to reliably differentiate synthetic wheat from common wheat as efficiently as the Infinium wheat 90K array is shown in Figure S3.

The final category included linked SNP for key breeding traits and SNP linking major germplasm resources genotyped with different technologies. In total, 457 wheat and 178 barley SNP corresponded to published trait-linked markers with 109 SNP associated with agronomically important genes (Table S1). Another 614 SNP provide a direct link to 19,778 GBS genotyped domesticated barley accessions (Milner *et al*. 2019).

### Assay performance – Single sample hybridsations

A limitation of hybridisation-based genotyping arrays is that their oligonucleotide probes hybridise both to the targeted locus and its homoeologues and paralogues if present (Cavanagh *et al*. 2013; Wang *et al*. 2014). Consequently, the ratio of allele-specific fluorescent signals observed for an assay depends on the locus copy number in the genome, with increasing copy number reducing the allele-specific fluorescent signal ratio and separation of SNP allele clusters. Further, SNP assay scorability and genotype calling can be confounded by the presence of mutations that modify oligonucleotide annealing such that different cluster patterns are observed across germplasm (Wang *et al*. 2014). An ideal assay design for a hybridisation-based genotyping array is therefore an oligonucleotide probe that binds at only one locus in the genome and has no known nucleotide variation underlying the probe hybridisation site. Theoretically this should ensure three distinct clusters corresponding to the genotypic states (REF, HET and ALT) expected of a single copy biallelic SNP. The increasing availability of genomic resources is now allowing this historical problem to be addressed. Hence, we used the combination of reference genome assemblies and genotypic data for large globally diverse wheat and barley collections to specifically target the design of single copy biallelic SNP assays.

For the purpose of evaluating the performance of the array, the wheat and barley diversity populations were used to define cluster positions for SNP genotype calling. The vast majority (98%) of the 39,654 SNP assays on the array produced scorable cluster patterns when hybridised with a barley or wheat sample; 91% (12,949/14,261) of the barley and 83% (20,090/24,598) of the wheat SNP assays could be reliably scored as single-copy biallelic markers, with the REF and ALT clusters having Theta values close to 0 and 1 in GenomeStudio SNP plots (Figure 5). While the remaining SNP could typically be reliably scored as biallelic markers, they showed cluster compression indicative of multiple loci. Few assays showed complex clustering patterns indicating the success of designing probes without underlying polymorphism. Five and 7% of wheat and barley assays showed a clustering pattern typical for the presence of a null allele. The occurrence of assays not behaving as single-copy biallelic markers reflects current knowledge gaps for structural variation in the genomes of wheat and barley including both copy number variation and presence-absence variation (Wang *et al*. 2014, Balfouier *et al*. 2019, Walkowiak *et al*. 2020).

**Figure 5.**
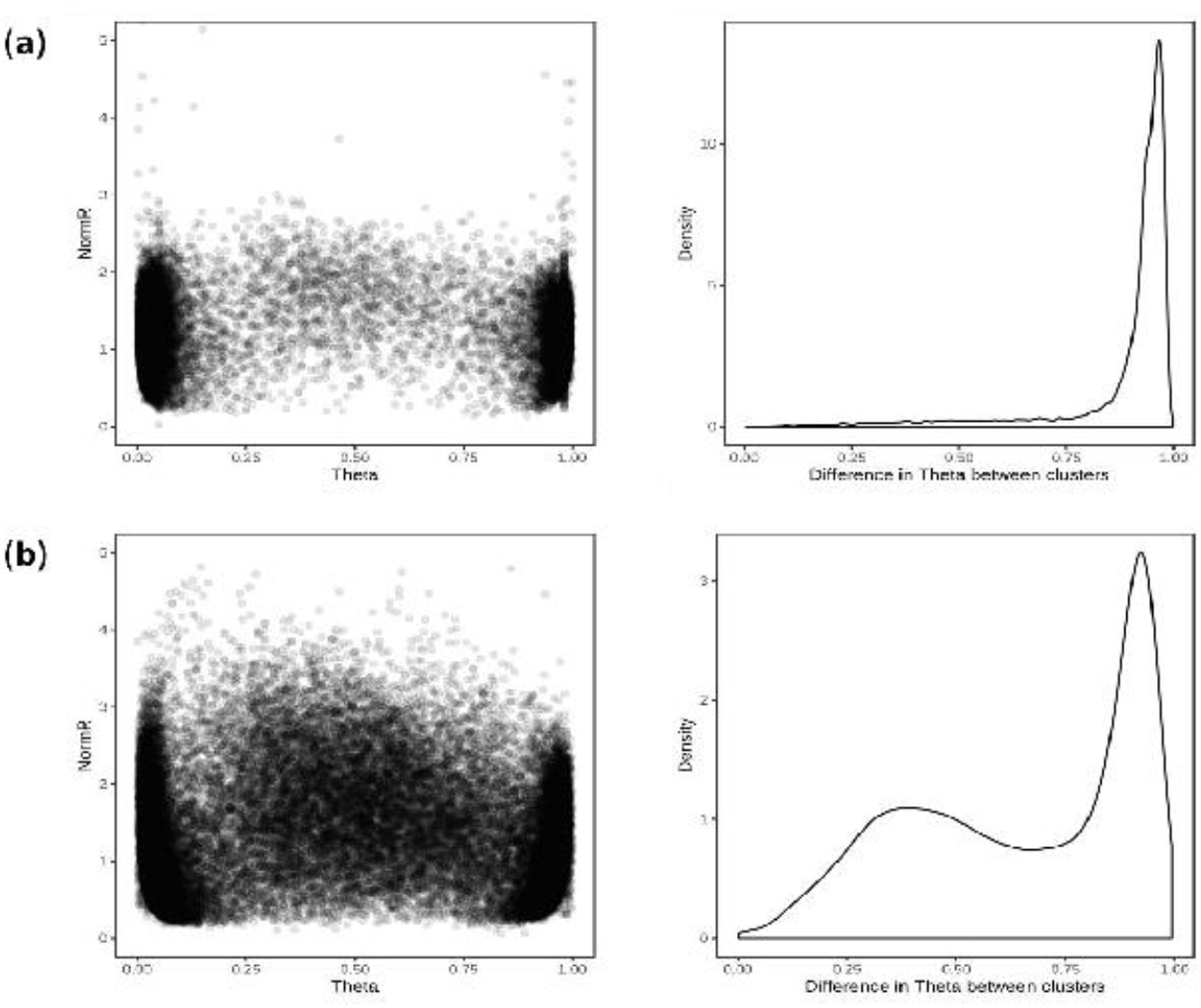
Cluster positions and theta separation of SNP in single sample hybridisation assays. Scatter plot of cluster positions (**left**) and density plot of difference in theta value between REF and ALT clusters (**right**) for (**a**) 14,261 barley and (**b**) 24,598 wheat SNP revealing polymorphism in the globally diverse wheat and barley populations

The concordance between called and actual genotypes was exceptionally high for both wheat and barley. The genotype concordance and correlation were 99.5 and 98.1% respectively in wheat when heterozygous genotype calls were excluded, and 97.6 and 95.7% when heterozygous calls were included. Similarly, 99.8% concordance and 99.2% correlation were observed in barley when heterozygous calls were excluded, and 98.2 and 97.2% was observed with heterozygous calls included. The average missing data rates was 4.8 and 3.8% in wheat and barley, respectively.

### Assay performance – Dual sample hybridisations

The design process specifically aimed to select species-specific SNP probes and thus it should be theoretically possible to jointly hybridise a wheat and barley sample to the same bead chip array (dual hybridisation) without loss of genotype calling accuracy. Cross-hybridisation between species is expected to confound genotype calling accuracy by creating shifts in SNP cluster positions and/or complex clustering patterns that cannot be easily scored.

To evaluate assay performance of a dual hybridisation, samples from the InterGrain commercial barley and wheat breeding programs were used to define cluster positions and call SNP genotypes for 576 dual hybridisation assays. The same samples were also assayed in single sample hybridisation assays to enable genotype calling accuracy between dual and single hybridisation assays to be directly compared.

Most of the barley and wheat SNP in dual hybridisation assays produced scorable cluster patterns. Shifts in cluster positions were observed, which indicated either that some oligonucleotide probes showed a degree of cross-species hybridisation or that deviation from the standard amount of sample DNA (200 ng per sample) recommended for the bead chip assay affected signal-to-noise. Through empirical testing, we found the quantity of genomic DNA per sample was a major factor causing shifts in cluster position (data not shown) and could be minimised by adjusting the input DNA for each sample to match the ratio of the genome size for each species; e.g. 200 ng barley DNA and 600 ng wheat DNA; the bread wheat genome is about three times larger than that of barley.

For the purpose of assessing genotype calling accuracy for dual hybridisation assays, only SNP that revealed polymorphism among the 576 wheat and barley samples assayed were considered. Of the 9,826 barley and 9,118 wheat SNP showing polymorphism, the vast majority were easily scored as biallelic markers and had good cluster separation, indicating that oligonucleotide probe cross-species hybridisation was minimal (Figure 6). The average concordance between genotypes calls for the same wheat and barley samples in single and dual sample hybridisation assays was 99.9, 96.7, and 99.8% for the REF, HET and ALT alleles, respectively. The average missing data rate across the wheat and barley samples was similar for both assay types, with 4.7 and 2.0% in dual and single hybridisation assays, respectively.

**Figure 6.**
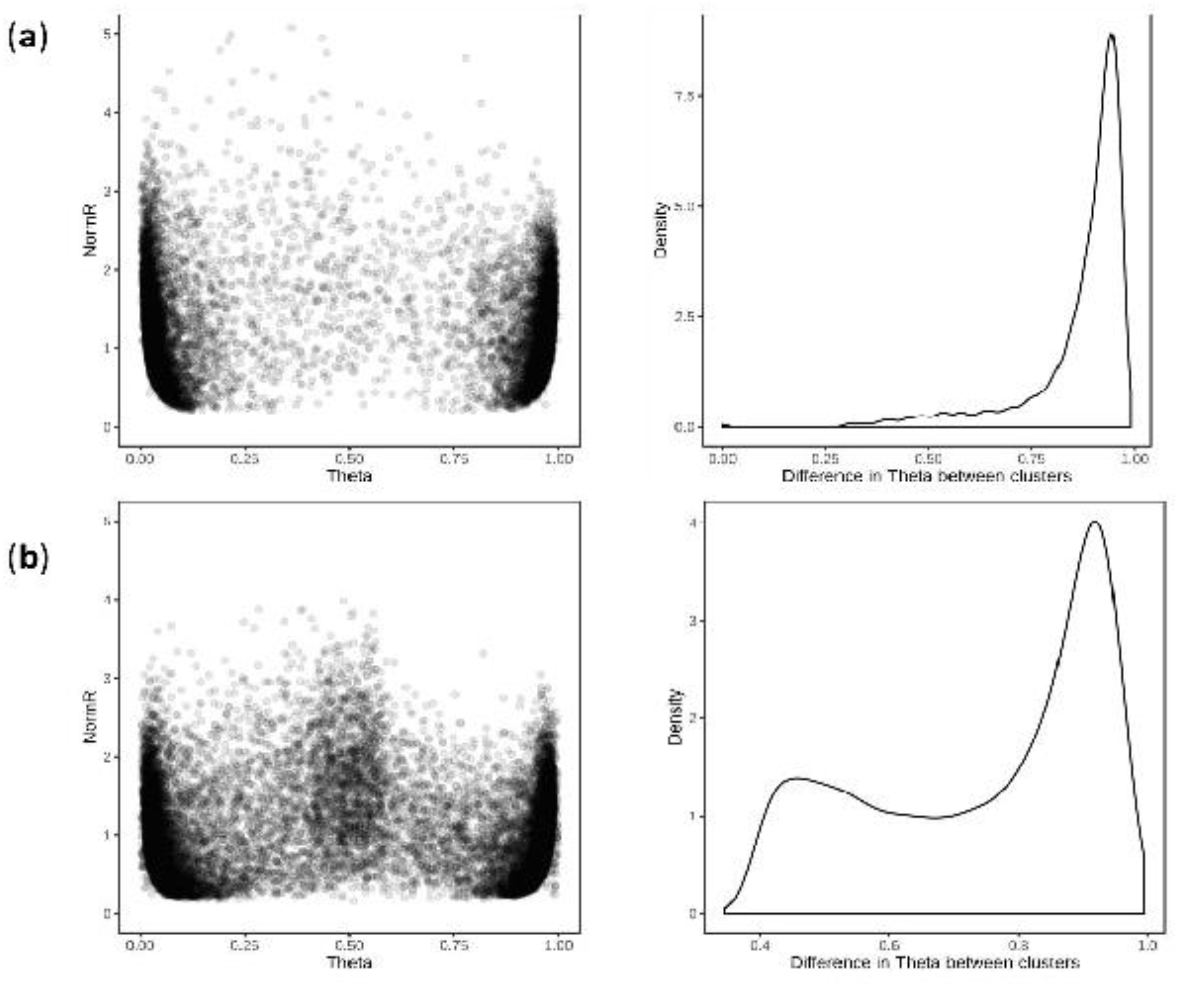
Cluster positions and theta separation of SNPs in dual hybridisation assays. Scatter plot of cluster positions (**left**) and density plot of difference in theta value between REF and ALT clusters (**right**) for (**a**) 9,826 barley and (**b**) 9,118 wheat SNP revealing polymorphism among 576 wheat and barley breeding lines

## Discussion

High-throughput, low-cost and flexible genotyping platforms are required for both research and breeding applications. Compared to GBS and PCR-based marker systems, array-based genotyping platforms are highly commercialised and highly customisable, both for the number of markers and samples assayed. They also have low genotype error and missing data rates compared to GBS technologies (Rasheed *et al*. 2017). Consequently, SNP arrays are widely utilised and several low-density SNP genotyping arrays have been developed for wheat and barley. Here, we described a novel approach that is applicable to any animal or plant species for the design of cost-effective, imputation-based SNP genotyping arrays with broad utility and that support the hybridisation of multiple samples to the same SNP array. The utility of the approach was demonstrated through the development of the Infinium Wheat Barley 40K SNP array.

The key difference between Infinium Wheat Barley 40K SNP array and previously reported array-based genotyping assays is a paradigm shift in the logic underpinning its design. To date, commonly used low-density genotyping arrays are comprised of the most scorable and informative markers from higher density arrays. For example, the Infinium Wheat 15K SNP array (Soleimani *et al*. 2020) and Axiom Wheat Breeders’ 35K SNP array (Allen *et al*. 2015) are derived from the Infinium Wheat 90K SNP array (Wang *et al.* 2014) and Axiom Wheat 820K SNP array (Winfield *et al.*, 2016). SNP on the Infinium 90K SNP array were derived from transcriptome sequence of 26 bread wheat accessions, while those on the Axiom 820K array were based on exome capture sequence from 43 bread wheat and wild species accessions representing the primary, secondary and tertiary gene pools. While these derived low-density arrays are affordable for routine deployment in breeding and research, their content is breeder-oriented and has limited utility outside the primary gene pool of hexaploid wheat.

The design implemented in the Infinium Wheat Barley 40K SNP array is based on the hugely expanded genotypic and genomic resources now available for wheat and barley. By using these resources, we were able to identify species-specific single-copy tSNP that capture a large proportion of the haplotypic diversity in globally diverse germplasm, are highly scorable for accurate genotype calling, minimise ascertainment bias and enable accurate imputation to high SNP density. In the case of wheat, this included the use of 2.04M SNP identified from exome sequence data of 1,041 accessions selected to maximally capture genetic diversity among a global collection of 6,700 accessions genotyped using the Infinium 90K SNP array (He *et al.* 2019; Figure 1a). The global collection included landraces, released varieties, synthetic derivatives, and novel trait donor and historical breeding lines. For barley, this included 932,098 SNP identified from exome sequence data of 267 accessions selected to maximally capture geographic diversity among landraces (Russell *et al*. 2016; Figure 1b), as well as SNP identified from target capture sequencing of 174 flowering time-related genes performed in 895 worldwide accessions (Hill *et al.* 2019). The latter dataset included global diverse cultivated and landrace germplasm.

By selecting tSNP enabling accurate imputation of common haplotype block diversity in globally diverse germplasm, the Infinium Wheat Barley 40K array is expected to maintain power for GWAS, genetic mapping and genomic selection (Jordan *et al*. 2015, He *et al*. 2015, Negro *et al*. 2019, Nyine *et al.* 2019). Haplotype blocks are essentially fixed stretches of DNA sequence that show little historical evidence of recombination and are effectively inherited as genetic units that are shuffled and assembled during breeding. The univariate LD metric *r*^2^ has been used in many tSNP algorithms as it is a major determinant of imputation accuracy and has a simple inverse relationship with the sample size required to detect associations in GWAS (Carlson *et al.* 2004, Ding and Kullo 2007). By selecting tSNP with an *r*^2^ ≥0.9 cut-off, we aimed to retain most of the information content in the original SNP set and to balance the power loss with the effort needed to compensate with increased sample numbers in downstream GWAS (~11%; i.e. 1/0.9). A significant advantage for using *r*^2^ is that it allows a high degree of flexibility in the composition of the final tSNP set, thereby enabling other design criteria to be applied without compromising overall tagging efficiency. This was especially important for implementing array design principles such as for selecting species-specific single-copy SNP targets that had no nucleotide variation underlying the probes to both maximise SNP scorability and support dual sample hybridisation assays. The success of our approach was confirmed by >97% accuracy (as measured by both correlation and concordance between the imputed and actual SNP genotypes) for imputing the set of SNP tagged at *r*^2^ ≥0.9 (inclusive of heterozygous calls) in both wheat and barley. Importantly, imputation accuracy was also high for the set of SNP tagged at *r*^2^ ≥0.5 (Table 1). To futureproof the array design, we added tSNP tagging genomic regions in wheat and barley that had sparse exome SNP coverage but high LD. We expect this content will similarly support accurate imputation to whole genome sequence once genomic resources needed to achieve this are available.

In emphasising the design focus on selecting tSNP for imputation, we also point out the limitations it has for fully capturing haplotype diversity in global wheat and barley germplasm. First, we did not tag LD blocks comprised of fewer than 10 SNP since this would have required an order of magnitude more SNP assays on the array; about 30,000 tSNP per species was required to tag about half of the non-singleton exome SNP at *r*^2^ ≥0.9 in each of wheat and barley (Figure 2). This presents a limitation for trait mapping using GWAS (but not genetic mapping) since trait loci located in untagged LD blocks will become increasingly harder to detect as their LD with a SNP on the array decreases. This limitation can be partly overcome by increasing sample size but is an unavoidable consequence of low-density arrays, despite our tSNP selection algorithm ensuring that we maximised the number of SNP tagged in LD. And second, in wheat the set of SNPs and LD relationships between them is still limited by the data currently available. As exome capture sequencing assays only 2-3% of the genome, the SNP discovered represent just a fraction of the true SNP density. It is therefore possible that SNP were not selected simply because the haplotype they represent was only sampled by a small number of SNP in that region and was below our selection thresholds. This limitation will only be overcome by large-scale whole genome sequencing efforts which are just beginning to become affordable for large genome-sized species. It should be noted that the LD patterns detected in this study will remain valid even with higher density sequencing and that the majority of the tagged LD haplotypes span across capture regions and so the number of SNP in high LD with the selected tSNP will only increase as higher density SNP data becomes available.

An argued advantage for GBS assays is that they are ascertainment bias free. Ascertainment bias can result in rare alleles being missed and genetic diversity being underestimated in non-ascertained populations (Clark *et al*. 2005), with its impact dependent on the study being undertaken. Increasing marker density and including low MAF markers in GWAS boosts power for QTL detection (Negro *et al*. 2019, Fikere e*t al.* 2020). Chu *et al*. (2020) reported that very low frequency markers (MAF <0.05) contributed to an improvement of genomic prediction accuracy in 378 winter bread wheat genotypes, and combined with the expectation that valuable novel diversity is most likely rare (Mascher *et al*. 2019), suggests that rare markers deserve careful consideration. Our tSNP selection algorithm prioritises haplotypes that diverge significantly from the reference genome used for SNP discovery in order to maximise the number of SNP tagged in LD; it is agnostic to the MAF of individual SNP (beyond the MAF cut-offs of 1% and 5% in wheat and barley, respectively). Consequently, the MAF spectrum of the wheat and barley tSNP closely resembled that observed for both the sets of tagged SNP and the filtered SNP in the globally diverse collections (Figure 3). Hence, we suggest the Infinium Wheat Barley 40K array has minimal ascertainment bias. Since tagging all minor variants is not feasible using low-density arrays, a better solution is to add minor variants into future versions of the array as trait associations are discovered, essentially as we have currently done for published trait linked markers.

To drive efficiencies for large-scale genotyping in commercial breeding programs, we explored the limits of the Infinium bead chip technology. One advantage of this technology is that each oligonucleotide assay probe has a unique physical position on the bead chip. This allows SNP arrays to be designed to genotype multiple crop species, with a user-defined number of SNP assigned to each species. The Infinium Wheat Barley 40K array assays 25,393 SNP in wheat and 14,261 SNP in barley. To the best of our knowledge, multispecies SNP arrays have only been used to assay a single sample at the time. Here, we demonstrated that through careful selection of species-specific oligonucleotide probes it is possible to jointly hybridise a wheat and barley sample to the same bead chip array, without substantial loss of genotype calling accuracy (Figure 6). The selection of such probes is facilitated by our design concept which exploits LD to identify SNP that can be considered equivalent for the purpose of genotyping. From a deployment perspective in a commercial breeding program, dual hybridisation doubles genotyping throughput, since twice as many samples can be processed given the same amount of time and resource. Dual hybridisation genotyping is potentially a game changing option for the adoption of genomics technologies by breeding companies that have large numbers of samples that can be co-ordinated into genotyping.

To ensure broad utility in research and breeding, we added SNP content capturing genetic diversity in the secondary and tertiary gene pools of wheat. This included 2,609 SNP from the Infinium 90K SNP array (Wang *et al*. 2014) tagging tetraploid wheat (A- and B-genome) diversity and clearly delineating tetraploid wheat from other types of wheat, as well as tetraploid species and subgroups from one another. Each SNP is single copy in tetraploid wheat and has been genetically and physically mapped (Maccaferri *et al.* 2019). It also included 1,206 single-copy SNP tagging *Ae. tauschii* (D-genome) diversity represented in 100 primary synthetic wheats, where each primary synthetic was derived from a different *Ae. tauschii* accession. Collectively, these SNP provide broad utility ranging from the differentiation and genetic characterisation of tetraploid and synthetic wheat (as well as other secondary and tertiary gene pools of wheat) to the tracking of introgressed genomic segments during breeding. Also included are SNP that directly link to the Infinium 90K (Wang *et al*. 2014) and 15K (Soleimani *et al*. 2020) wheat arrays to ensure connectivity with legacy genotypic datasets and research. For barley, we included 685 SNP that overlap with SNP reported for 19,778 GBS genotyped accessions from the IPK Genebank (Milner *et al*. 2019) to provide a direct anchor to that resource, and 1,239 SNP that overlap with the Infinium 50K barley SNP array (Bayer *et al*. 2017) which link to 21,606 common SNP following imputation. Finally, we included trait-linked SNP and SNP tagging GWAS signals for key breeding and research targets reported in the published literature.

The overall array design makes it ideal for a wide range of research and breeding applications, from germplasm resource characterisation, GWAS and genetic mapping to tracking introgressions from different sources, marker-assisted breeding and genomic selection. Its utility is further enhanced through the web-based tool *Pretzel* (Keeble-Gagnère *et al.* 2019; https://plantinformatics.io/) which enables the array’s content to be visualised and interrogated in real-time in the context of numerous genetic and genomic resources. For example, the SNP can be visualised relative to the genetic and physical positions of other DNA marker types (e.g. SSRs, DArT), SNP on other genotyping arrays, trait loci, annotated genes and syntenic positions in the genomes of other crops and model species. The ability to upload and visualise data in *Pretzel* allows breeders and researchers to seamlessly link and interrogate their own data in the context of publicly available datasets hosted in *Pretzel*. Combined, the Infinium Wheat Barley 40K SNP array and *Pretzel* enable legacy and current research to seamlessly connect to breeding.

In conclusion, we have described a novel approach applicable to any animal or plant species for designing cost-effective imputation-enabled SNP genotyping arrays which have broad applicability in research and industry applications (e.g. GWAS, genomic prediction and operational breeding) and support the hybridisation of multiple samples to the same array. The utility of this design approach was demonstrated through its implementation to develop a new Infinium Wheat Barley 40K SNP array. In addition, to supporting broad utility in research and breeding, this array can be used as a resource to connect genetic and genomic datasets generated across germplasm pools and time. The array is further supported by the publicly available web-tool *Pretzel* and is available for purchase by the international wheat and barley community from Illumina Ltd, the manufacturer of the Infinium bead chip technology.

## Data Statement

Exome data used from Russel *et al.* 2016 and He *et al.* 2019 are accessible under EBI ENA project accession numbers PRJEB8044 and PRJEB31218, respectively. The filtered set of exome genotype calls for accessions and SNP underpinning the LD analysis and tag SNP selection for wheat (https://doi.org/10.7910/DVN/5LVYI1) and barley (https://doi.org/10.7910/DVN/CUPAXD) as well as the D-genome synthetic derivative-enriched SNP matrix (https://doi.org/10.7910/DVN/0QEASF) are available through Dataverse at https://dataverse.harvard.edu/dataverse/WheatBarley40k_v1. Information about the status of each SNP, including tag SNP set ID and whether the SNP passed design filters, is included in the INFO column. Illumina 90k iSelect genotypes for the accessions used to select tetraploid-specific content is available at https://figshare.com/articles/dataset/Durum_Wheat_cv_Svevo_annotation/6984035 (Maccaferri *et al*. 2019).

## Author Contributions

R.P. performed LD analysis. G.K-G selected tagging SNP, performed imputation analyses and produced the final designs. K.F. and D.W. performed exome and whole genome sequencing, Infinium Wheat Barley 40K assays and genotype calling. J.T. performed sequence alignments and genotype calling. H.R., J.G., A.R., D.M., D.M., selected non-tagging SNP and provided wheat and barley germplasm. T.W., H.D, J.T. and M.H. conceived the project. G.K-G and M.H. wrote the manuscript.

## Conflict of Interest

The authors declare that they have no conflict of interest.

## Supporting Information

**Table S1**. Detailed description of Infinium Wheat Barley 40K SNP array content

**Figure S1**. Cumulative number of SNP tagged by tSNP at *r*^2^ ≥0.90 in each chromosome in wheat and barley. Curves shown until the first singleton SNP is reached on each chromosome

**Figure S2**. PCA based on (**a**) 17,600 SNP described in Maccaferri *et al.* (2019) from the Infinium wheat 90K SNP array and (**b**) 2,609 SNP selected for inclusion on the Infinium Wheat Barley 40K SNP array showing differentiation among 1,856 tetraploid wheat accessions representing wild emmer wheat from North Eastern Fertile Crescent (WEW-NE), wild emmer wheat from Southern Levant Fertile Crescent (WEW-SL), domesticated emmer wheat (DEW), domesticated emmer wheat from Ethiopia (DEW-ETH), durum wheat landraces (DWL) and durum wheat cultivars (DWC)

**Figure S3**. PCA based on (**a**) 37,105 called SNP from the Infinium wheat 90K SNP array, and (**b**) 20,665 SNP on the Infinium Wheat Barley 40K SNP array showing differentiation among bread wheat (green), synthetics derivatives (blue) and hexaploid wheat derived from crosses between bread and durum accessions (red) (number of accessions=1219)

## Notes

### Competing Interest Statement

The authors have declared no competing interest.

