## Supplementary figures and images for "Novel design of imputation-enabled SNP arrays for breeding and research applications supporting multi-species hybridisation"

### Figure S1

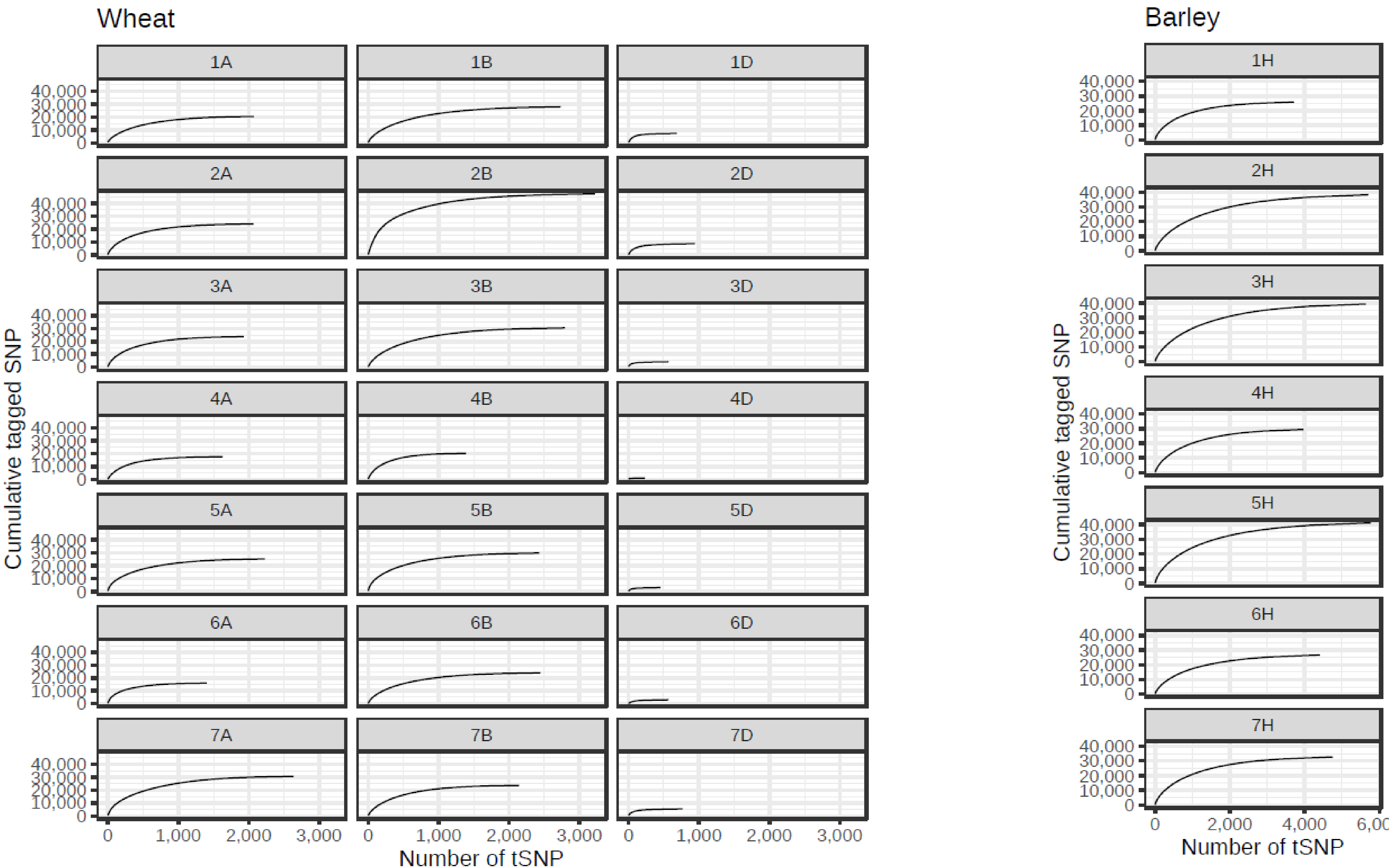

### Figure S2

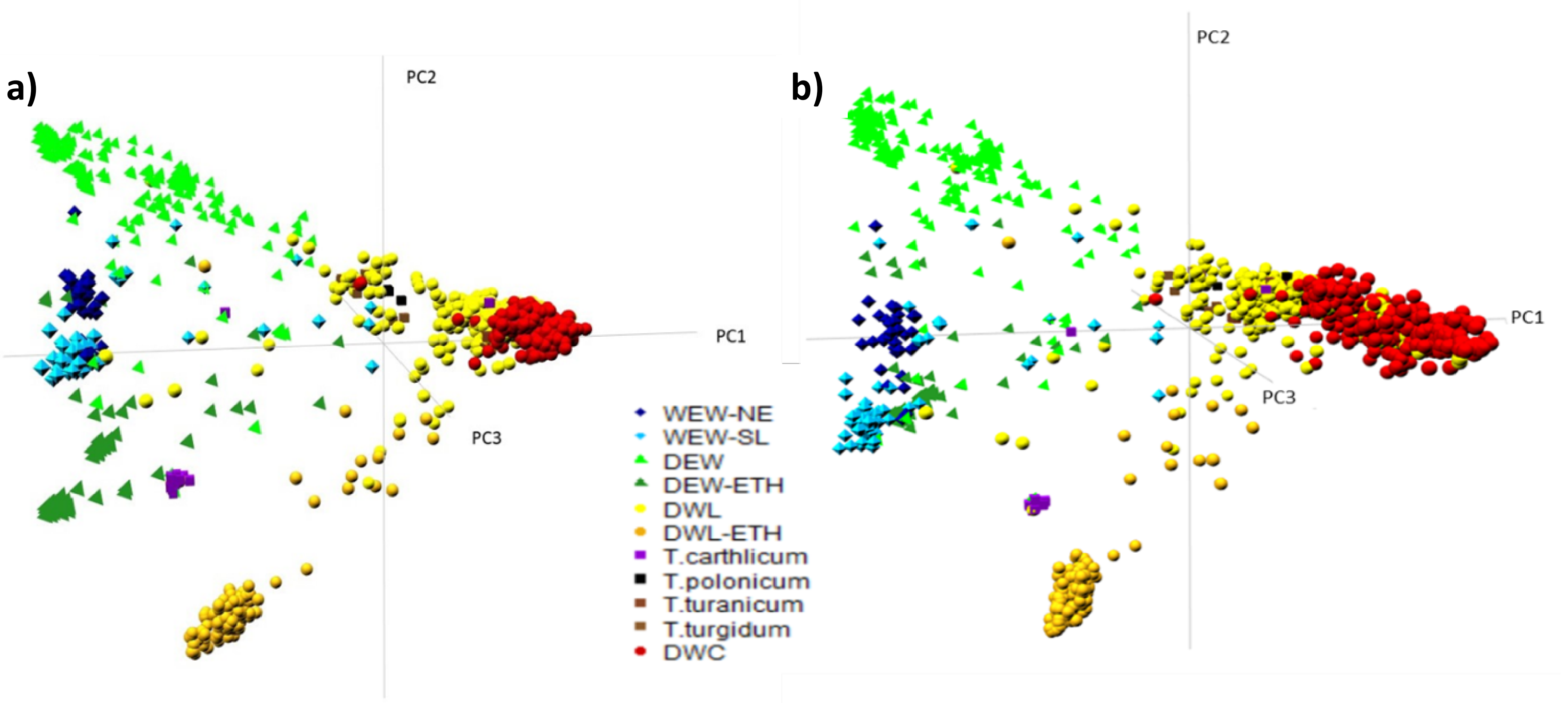

### Figure S3

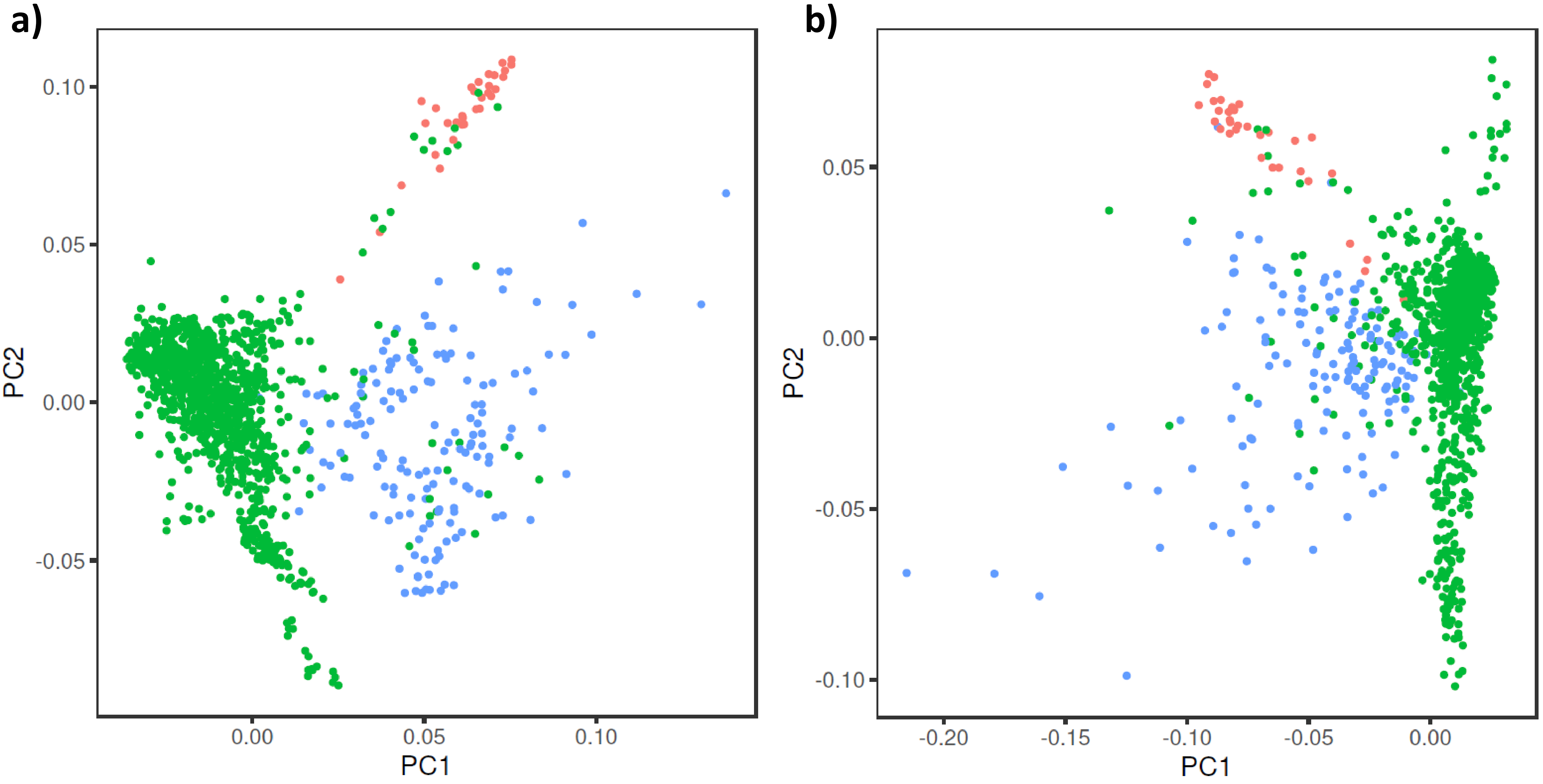
